# The Vesicular Glutamate Transporter Modulates Sex and Region-Specific Differences in Dopaminergic Neuron α-Synuclein Toxicity by Modifying Cytosolic Dopamine Levels

**DOI:** 10.64898/2026.01.17.699798

**Authors:** Kevin Garzillo, Mia Perulli, Daniel T Babcock

## Abstract

Parkinson’s disease disproportionately affects males; however, the cause of this sex difference is unknown. We found that expressing mutant α-synuclein ^A53T^ in *Drosophila* dopamine neurons recapitulates the sex differences observed in human Parkinson’s disease patients. Male flies exhibited greater age-related motor impairment and more severe dopamine neuron degeneration than females. Selective masculinization of female dopamine neurons via knockdown of the sex determination gene *Transformer* eliminated the observed sex differences in locomotor ability and neurodegeneration by increasing the severity of motor defects and degeneration in females. *Transformer* knockdown in dopamine neurons also reduced total vesicular glutamate transporter staining in the brain. Direct knockdown of the vesicular glutamate transporter in female dopamine neurons expressing α-synuclein ^A53T^ exacerbated motor dysfunction, altered mitochondrial dynamics, and accelerated dopamine neuron degeneration. Increasing cytosolic dopamine via knockdown of the vesicular monoamine transporter or increasing total dopamine levels via levodopa treatment phenocopied vesicular glutamate transporter knockdown; furthermore, reducing total dopamine via alpha-methyl-p-tyrosine treatment protected against vesicular glutamate transporter knockdown. These results support a model in which lower VGLUT levels in dopamine neurons result in higher levels of cytosolic dopamine, which leads to dopamine mediated mitochondrial dysfunction and increased susceptibility to α-synuclein ^A53T^ toxicity.

## Introduction

Parkinson’s disease (PD) is the fastest growing neurodegenerative disease worldwide^1^. Its canonical symptoms include bradykinesia, tremor, rigidity, and accumulation of Lewy bodies ^2^. The motor symptoms of PD arise due to the progressive loss of dopamine (DA) neurons in the substantia nigra pars compacta (SNc) ^2^. There are no disease modifying therapies for PD ^3^. Current treatment is centered around symptom management with the mainline approach being administration of levodopa (L-DOPA), either as monotherapy or in combination with DA agonists or monoamine oxidase inhibitors^3^. PD disproportionately affects males, as females are less likely to develop PD, tend to manifest motor symptoms later in the disease, and self-report less severe motor impairments ^4–8^. It is unknown whether these sex differences are due to intrinsic differences in biology between males and females or differences in environmental exposure to PD linked toxicants; however, data from multiple animal models supports the former conclusion, as males in these studies tend to exhibit more severe neurodegeneration than females ^9^.

Most PD cases are considered idiopathic ^2^. Despite this incomplete understanding of PD etiology, multiple genetic variants have been linked to PD, most notably the A53T mutation in the gene *SNCA* ^2,10^. *SNCA* encodes α-synuclein (αSyn), a 140 amino acid synaptic protein that is the primary component of Lewy bodies ^11^. αSyn ^A53T^ causes PD via a toxic gain of function, as the A53T mutation is autosomal dominant and expression of αSyn ^A53T^ in model organisms leads to neurodegeneration ^10,12–16^. Multiple reports suggest that αSyn ^A53T^ interacts with mitochondria, altering ATP production, disrupting fission/fusion dynamics, and modifying mitophagy, which leads to a dysfunctional mitochondrial network and increased production of reactive oxygen species (ROS) ^17–23^. DA neurons are particularly vulnerable to oxidative stress, as cytosolic DA can undergo spontaneous auto-oxidation, forming both ROS and toxic dopamine quinones (DAQs); furthermore, targeted degradation of cytosolic DA via monoamine oxidases (MAOs) also generates ROS as well as 3,4-dihydroxyphenylacetaldehyde (DOPAL), a toxic DA metabolite ^24–29^. Oxidative environments increase αSyn’s propensity to form aggregates ^30–34^, and αSyn aggregates preferentially interact with mitochondria, further impairing mitochondrial function and exacerbating oxidative stress ^21,35–37^. Impaired mitochondria produce more ROS, creating a positive feedback loop that amplifies oxidative stress and αSyn aggregation, terminating in apoptosis ^38,39^. Therefore, the selective degeneration of DA neurons in PD may be due to their high burden of basal ROS and reactive DA species, which stress mitochondria and increase their susceptibility to further damage. This model is supported by postmortem analysis demonstrating that PD patients have elevated levels of midbrain ROS and mitochondrial mutations ^40–42^. However, not all midbrain DA neurons are equally vulnerable to αSyn pathology. In the midbrain of PD patients, both SNc and ventral tegmental area (VTA) DA neurons exhibit degeneration but a higher percentage of SNc DA neurons are lost than VTA DA neurons ^43–45^. This trend of increased vulnerability of SNc DA neurons relative to VTA DA neurons has been recapitulated in rats expressing human αSyn ^A53T^ ^46^. In addition to this variability in degeneration between regions, vulnerability varies within regions during disease progression, with some DA neurons dying early, others late, and some not at all ^43^. The mechanism underlying this selective vulnerability remains unknown and is an ongoing area of research.

These differences in vulnerability between individual neurons, brain regions, and sexes may be due to differential expression of the vesicular glutamate transporter 2 (VGLUT2). A subset of DA neurons express VGLUT2 and co-release both DA and glutamate ^47–49^. In both animal models of PD and in postmortem human brain samples from PD patients, VGLUT2 expressing DA neurons were more resistant to degeneration than their non VGLUT2 expressing counterparts ^50,51^. Interestingly, in humans, mice, and fruit flies, expression of VGLUT2/VGLUT (the *Drosophila* VGLUT2 homolog) is higher in female DA neurons than in male DA neurons ^48^. Furthermore, the VTA has a larger percentage of VGLUT2 expressing DA neurons than the SNc ^51^. These facts taken together suggest that differences in VGLUT2 expression may in part account for region specific and sex specific differences observed in PD. However, the mechanism underlying this putative neuroprotection has not been determined. In DA neurons, VGLUT2 mediated storage of glutamate into synaptic vesicles augments vesicle acidification ^52^. This decrease in vesicle pH amplifies vesicular loading of DA by the monoamine/proton antiporter Vesicular Monoamine Transporter 2 (VMAT2) ^52,53^. Thus, it has been hypothesized that VGLUT2 confers resilience to DA neurons in PD by promoting vesicular sequestration of cytosolic, reactive DA; however, this proposed mechanism has not yet been experimentally tested ^9,47,51,53,54^.

Here, we demonstrate that sex and regional differences in αSyn ^A53T^ toxicity are due to differences in DA neuron VGLUT expression. Then we show that increasing cytosolic or total DA phenocopies VGLUT knockdown. Lastly, we show that VGLUT knockdown can be partially rescued by reducing total DA levels. These results support a model in which lower VGLUT levels in DA neurons result in higher levels of cytosolic DA, which leads to DA mediated mitochondrial dysfunction and increased susceptibility to αSyn ^A53T^ toxicity.

## Methods

### Drosophila stocks and husbandry

Fly stocks were housed at 25°C on standard *Drosophila* media. Experimental flies were collected after eclosion and separated by sex into groups of 10. Flies were then aged for the indicated number of days on standard *Drosophila* media at 25°C. During aging, they were transferred to vials containing fresh food every two days. For experiments with L-DOPA and α-methyl-p-tyrosine (AMPT), drugs were dissolved directly in *Drosophila* media at the indicated concentration. The following stocks were obtained from Bloomington stock center: UAS-VGLUT ^55^, UAS-VGLUT RNAi ^56^, UAS-VMAT RNAi ^56^, UAS-Tra RNAi ^56^, TH-GAL4 ^57^, UAS-Luciferase RNAi ^56^, UAS-MitoTimer ^58^, DVGLUT-GAL4 ^59^, and BDSC-US-N(#99772). 10XUAS-IVS-Syn21-GFP-p10 (JFRC81) was a gift from Gerald Rubin.

### Generation of HA tagged UAS-*α*Syn ^A53T^

UAS-HA: αSyn ^A53T^ was generated by inserting human αSyn ^A53T^ cDNA into an entry vector using the pCR8 Gateway cloning kit (ThermoFisher) and cloning into pTHW (DGRC Stock 1099), which includes the UASt promoter and an N-terminal 3XHA tag. The construct was inserted into the genome by BestGene Inc (Chino Hills, CA).

### Locomotor assay

Flies were transferred into 20 cm tall glass vials, each marked at 12 cm. Flies were allowed 60 seconds to acclimate and then the vial was tapped on a mouse pad to force the flies to the bottom of the vial. A successful climbing attempt was defined as a fly crossing the 12 cm mark within 15 seconds of being tapped down. This process was repeated twice for each group, and the best climbing attempt was recorded. Climbing Index is the percentage of successful climbing attempts per condition.

### Immunohistochemistry

Brains were extracted in PBS and fixed in 4% paraformaldehyde (PFA) for 40 minutes. Samples were then washed 5 times with PBST and incubated in blocking buffer (PBS, 0.1% goat serum, and 0.2% Triton X-100) for an hour. Samples were then incubated with primary antibodies for 24 hours. After incubation, samples were washed 5 times with PBST and then incubated with secondary antibodies for 2 hours. Samples were then washed 5 times with PBST and mounted in Vectashield (Vector Laboratories).

The following primary antibodies were used: rabbit anti-tyrosine hydroxylase (1:100, AB152, Millipore), rabbit anti-*Drosophila* VGLUT N-terminus (1:500, gift from Hermann Aberle ^60^), and chicken anti-GFP (ThermoFisher, #A10262). The following secondary antibodies were used: Alexa Fluor 488 goat anti-rabbit (1:200, Fisher Scientific), Alexa Fluor 568 goat anti-rabbit (1:200, Fisher Scientific), and Alexa Fluor 488 goat anti-chicken (1:200, Fisher Scientific)

### Confocal microscopy and fluorescence quantification

Brains were imaged using a Zeiss LSM 880 confocal microscope. PPL1, PPM1/2, and PPM3 cluster images were acquired using a 63x oil objective, and whole brain images were acquired using a 20x objective. Z stacks were formed into composites via ImageJ and brightness for each set of images was set using Adobe Photoshop.

For VGLUT fluorescence quantification, an ROI was drawn around the brain in ImageJ and mean fluorescence intensity was measured.

### Dopaminergic neuron quantification

Posterior DA neurons per cluster were quantified using a Nikon Eclipse Ni-U fluorescent microscope at a magnification of 20X. All slides were quantified blind with respect to sex, genotype, and drug treatment.

### Mitochondrial analysis

To quantify DA neuron mitochondrial number, morphology, and turnover we used MitoTimer, a genetically encoded, mitochondrially localized, modified DsRed that, when newly synthesized, exhibits a GFP-like fluorescence spectrum but irreversibly shifts to red upon oxidation ^58^. MitoTimer has been used in *Drosophila* to assay mitochondrial morphology and turnover ^58,76–79^. Since older mitochondria contain more oxidized MitoTimer, the red-to-green fluorescence ratio serves as a measure of mitochondrial age ^58^. To quantify DA neuron mitochondrial turnover, green and red fluorescence intensity for PPL1 and PB mitochondria z-stacks were measured via ImageJ. Red to green ratio was calculated by dividing the mean fluorescence of the red channel by the mean fluorescence of the green channel for each cluster. Mitochondria morphology for DA neuron clusters was analyzed using the Mitochondria Analyzer plugin for ImageJ with a block size of 1.45 and C-value of 5 ^61^.

### RNA extraction and RT-qPCR

3 days post eclosion, three groups of 20 heads per sex from TH-GAL4 > UAS-αSyn ^A53T^ flies were homogenized using a motorized pestle. RNA was extracted from homogenates using Monarch’s Spin RNA Isolation Kit (New England BioLabs). cDNA was synthesized from the extracted RNA using Invitrogen’s SuperScript IV VILO Master Mix. RT-qPCR was carried out using Applied Biosystems PowerUp SYBR Green Master Mix. Reactions were conducted in triplicate for each group and then averaged to obtain CT values for both αSyn ^A53T^ and Actin-5C. αSyn ^A53T^ primers: Forward 5′ AACCAAACAGGGTGTGGCAG 3′ and Reverse 5′ CCCTCCTTGGTTTTGGAGCC 3′. Actin-5C primers: Forward 5′ CGAAGAAGTTGCTGCTCTGGTTGT 3′ and Reverse 5′ GGACGTCCCACAATCGATGGGAAG 3′ ^62^. Relative αSyn ^A53T^ expression was calculated using the ΔΔCT method as previously described ^63^.

### Statistical analysis

Climbing, DA neuron number, fluorescence intensity, and mitochondrial morphology data were analyzed separately for each sex using one way ANOVA followed by Tukey’s post hoc test, except where otherwise noted. Analyses were designed to assess genotype/drug dependent effects within each sex, and relevant statistical comparisons are indicated in the figures. qPCR data was analyzed using a two tailed t-test. All statistical analysis was carried out using GraphPad Prism.

## Results

### Male but not female DA neurons are vulnerable to *α*Syn ^A53T^ pathology

Transgenic expression of αSyn ^A53T^ in *Drosophila* produces PD-like phenotypes, including an age-related decline in locomotor ability, degeneration of DA neurons, and intracellular inclusions ^15^. Two recent papers demonstrated that pan neuronal expression of αSyn ^A53T^ in *Drosophila* causes more severe locomotor defects in males than in females, recapitulating what is observed in humans ^64,65^. However, the molecular mechanisms and neuronal populations underpinning these sex differences are unknown. To determine if this sex specific difference in locomotor ability is due to sex differences in DA neuron vulnerability to αSyn ^A53T^, we used TH-GAL4 to express αSyn ^A53T^ specifically in DA neurons and then measured locomotor ability via the climbing assay at day 3 and day 35 post eclosion. DA neuron specific expression of αSyn ^A53T^ induced age dependent climbing defects in males but not females relative to both UAS-αSyn ^A53T^ /+ and TH-GAL4/+ (Figure 1A). Strong age dependent climbing defects are often indicative of neurodegeneration; therefore, to determine if αSyn ^A53T^ expression also induced sex specific DA neuron degeneration, we stained both male and female brains for tyrosine hydroxylase (TH), a marker of DA neurons. Males but not females expressing αSyn ^A53T^ had a reduction in TH positive cells in both the Protocerebral Posterior Lateral 1 (PPL1) and Protocerebral Posterior Medial 1 and 2 (PPM1/2) DA neuron clusters (Figure 1B-C). To confirm that the reduction was due to decreased cell number and not reduced TH signal, we selectively drove expression of either GFP alone or GFP with αSyn ^A53T^ in DA neurons and quantified the number of GFP positive PPL1 and PPM1/2 neurons in males and females. Consistent with our previous experiment, males but not females expressing αSyn ^A53T^ exhibited fewer GFP positive neurons in PPM1/2 compared to controls (Supplementary Figure 1A-B), confirming that the PPM1/2 DA neuron cluster is selectively vulnerable in males but not females to αSyn ^A53T^. Interestingly, expression of αSyn ^A53T^ for both males and females did not reduce the number of TH positive or GFP positive cells in the Protocerebral Posterior Medial 3 (PPM3) DA neuron cluster (Figure 1B-C & Supplementary Figure 1A-B), suggesting *Drosophila* exhibit both sex and region specific vulnerability to αSyn ^A53T^. Lastly, to ensure that differences in vulnerability between sexes were not due to differential transgene expression, we used RT-qPCR to measure mRNA levels in males and females expressing αSyn ^A53T^ and observed no differences in transgene expression (Supplementary Figure 2A).

**Figure 1:**
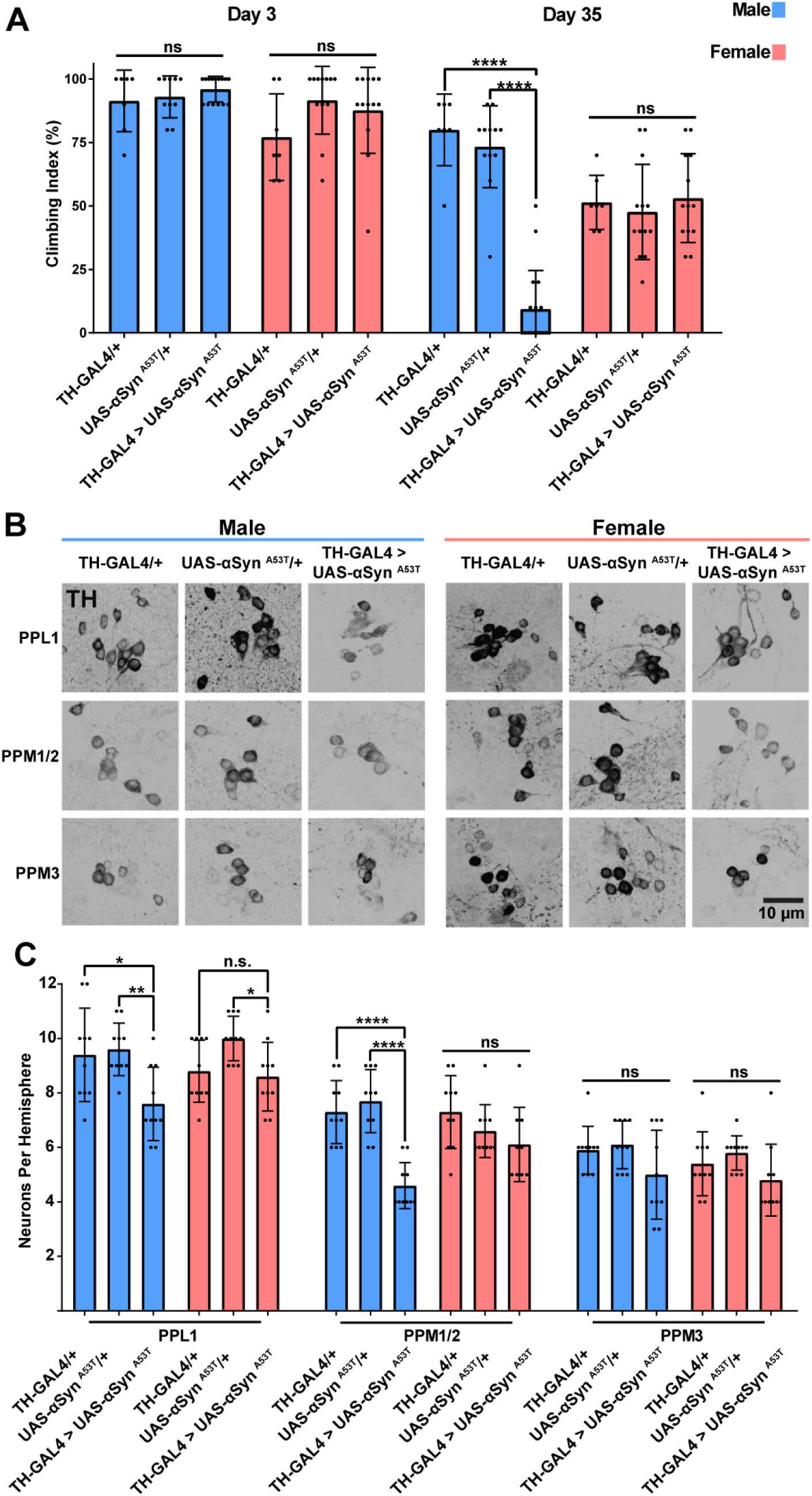
Male flies are selectively vulnerable to *α*Syn ^A53T^ toxicity. **A** Climbing assay data for male and female flies at 3- and 35-days post eclosion. Each point on the graph represents a vial of 10 flies. **B** Representative images of male and female PPL1, PPM1/2, and PPM3 DA neurons. TH immunofluorescence is black. **C** Quantification of male and female DA neurons 35 days post eclosion. For all graphs male data is blue and female data is pink. Error bars demonstrate standard deviation (SD). * < 0.05, ** < 0.01, *** < 0.001, **** < 0.0001, N.S = not significant.

### Sex differences in DA neuron vulnerability to *α*Syn ^A53T^ pathology are cell autonomous

Sex determination in *Drosophila* is mostly cell autonomous and is regulated by the RNA binding protein Transformer (*Tra*) ^66,67^. Previous reports have demonstrated that knockdown of *Tra* allows for selective masculinization of specific neuronal populations in females ^68–70^. Thus, to determine if sex specific differences in αSyn ^A53T^ vulnerability are due to cell autonomous factors, we selectively masculinized female DA neurons by expressing αSyn ^A53T^ with *Tra* RNAi or GFP (to control for transgene dilution) in DA neurons. Co-expression of αSyn ^A53T^ with *Tra* RNAi eliminated the sex differences in climbing ability (Supplementary Figure 3A) and DA neuron degeneration (Figure 2A-D) by increasing the severity of αSyn ^A53T^ induced climbing defects and degeneration of PPL1 and PPM1/2 DA neurons in females. Conversely, co-expression of αSyn ^A53T^ with *Tra* RNAi in male DA neurons did not modify the effects of αSyn ^A53T^ on climbing ability (Supplementary Figure 3A) or DA neuron degeneration (Figure 2A-F). These results taken together demonstrate that sex differences in DA neuron vulnerability to αSyn ^A53T^ are due to cell autonomous differences between males and females.

**Figure 2:**
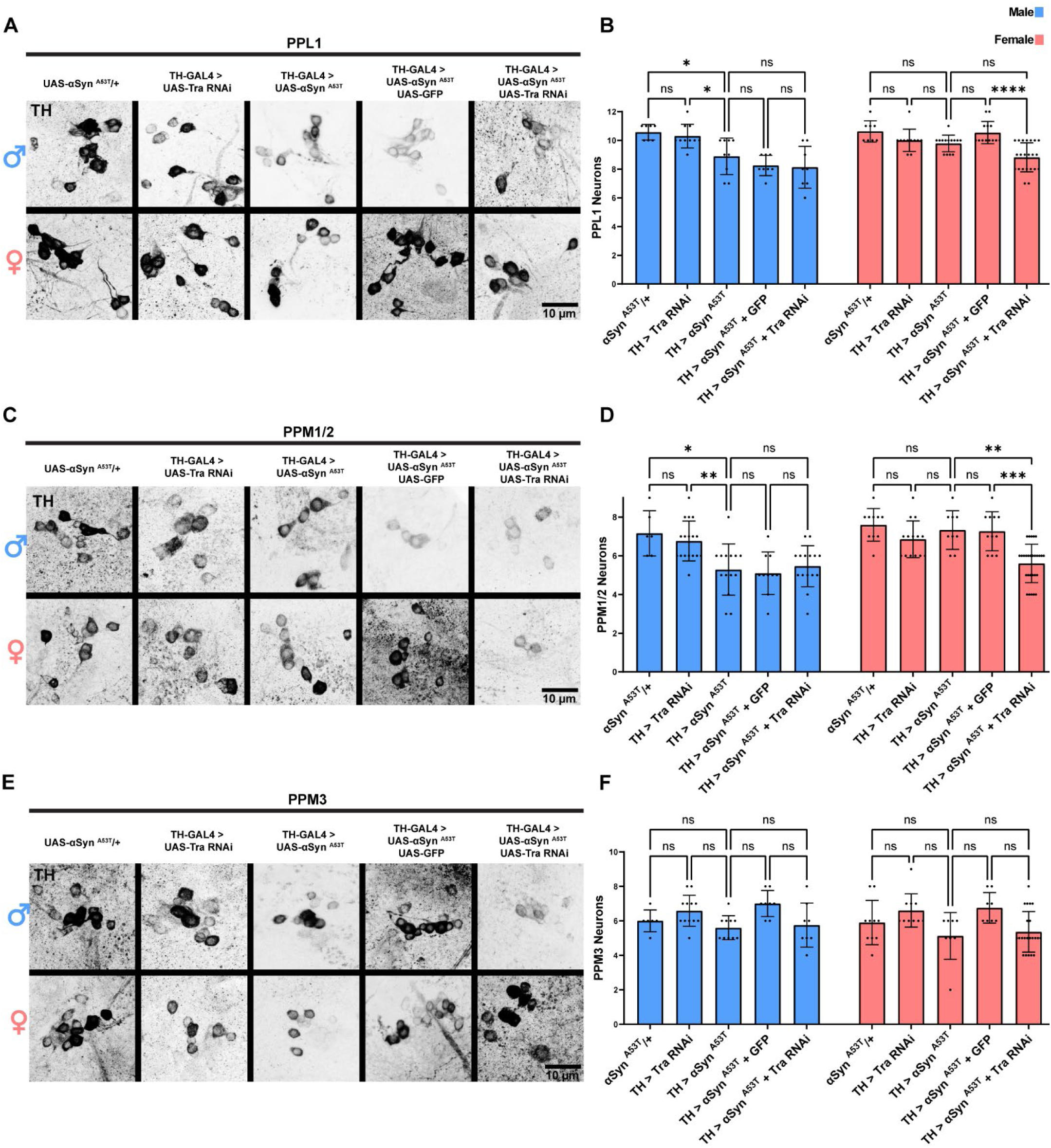
Sex differences in DA neuron vulnerability to *α*Syn ^A53T^ toxicity are cell autonomous. **A** Representative images of male and female PPL1 neurons 35 days post eclosion. **B** Quantification of male and female PPL1 neurons. **C** Representative images of male and female PPM1/2 neurons 35 days post eclosion. **D** Quantification of male and female PPM1/2 neurons. **E** Representative images of male and female PPM3 neurons 35 days post eclosion. **F** Quantification of male and female PPM3 neurons. TH immunofluorescence is black. For all graphs male data is blue and female data is pink. Error bars demonstrate SD. * < 0.05, ** < 0.01, *** < 0.001, **** < 0.0001, N.S = not significant.

### VGLUT knockdown in DA neurons abolishes sex and region-specific differences in DA neuron vulnerability to *α*Syn ^A53T^ pathology

Previous work has demonstrated that VGLUT expression is higher in female DA neurons than in male DA neurons and that reducing VGLUT levels can increase susceptibility to mitochondrial oxidative stress ^48,71,72^. Therefore, to determine if *Tra* modulates sex differences in VGLUT levels, we co-expressed αSyn ^A53T^ with *Tra* RNAi or GFP and then stained both male and female brains for VGLUT. αSyn ^A53T^ expression significantly reduced VGLUT staining in males but not females (Figure 3A-B). However, co-expression of αSyn ^A53T^ with *Tra* RNAi in females eliminated the sex difference by reducing total VGLUT staining in the brain. These results demonstrate that DA neuron Tra expression modulates sex differences in αSyn ^A53T^ induced changes to brain VGLUT levels.

**Figure 3:**
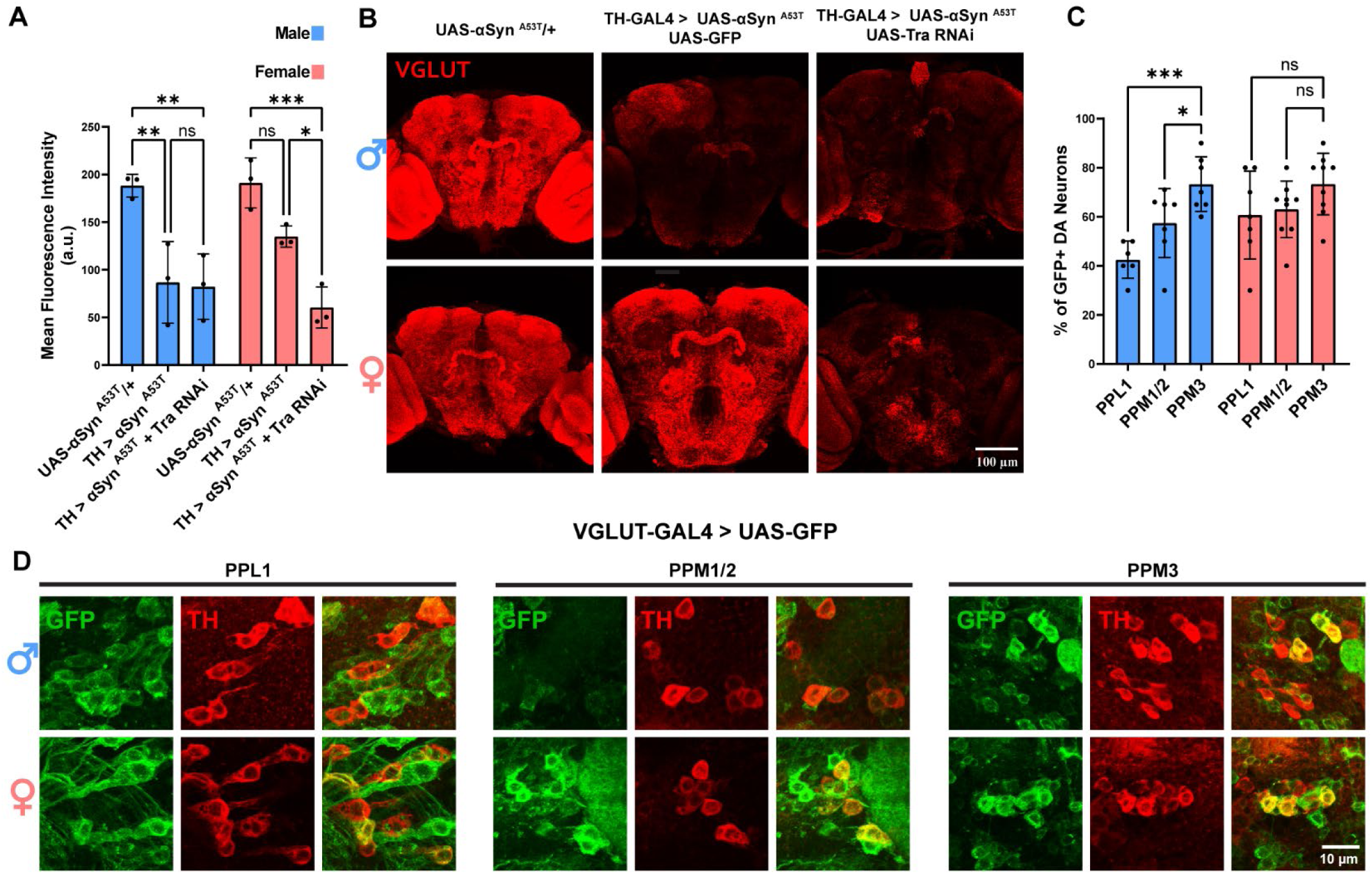
Tra expression modulates sex differences in *α*Syn ^A53T^ induced changes to brain VGLUT immunofluorescence. **A** Quantification of VGLUT immunofluorescence intensity. **B** Representative images of whole brain VGLUT staining 15 days post eclosion. VGLUT immunofluorescence is red. **C** Quantification for the percentage of GFP positive cells per DA cluster for males and females. **D** Representative images of VGLUT-GAL4 > UAS-GFP male and female PPL1, PPM1/2, and PPM3 DA neurons 15 days post eclosion. GFP and TH immunofluorescence are green and red, respectively. . Error bars demonstrate SD. * < 0.05, ** < 0.01, *** < 0.001, **** < 0.0001, N.S = not significant.

Masculinization of DA neurons increased vulnerability to αSyn ^A53T^ in PPL1 and PPM1/2 neurons, but not in PPM3 neurons. Therefore, we reasoned that if cluster specific differences in vulnerability are due to differences in VGLUT expression, then higher VGLUT levels and/or a greater percentage of VGLUT expressing neurons should be present in the PPM3 cluster relative to the PPL1 and PPM1/2 clusters in males. To test this, we used VGLUT-GAL4 to drive expression of GFP in VGLUT expressing neurons and then quantified the number of TH neurons that were GFP positive. GFP positive neurons were detected in all three clusters for both sexes; however, in males, but not females, the percentage of GFP positive DA neurons was significantly lower in PPL1 and PPM1/2 relative to PPM3 (Figure 3C-D). These results are consistent with the increased resilience of PPM3 neurons in males being mediated by VGLUT.

To determine if loss of VGLUT increases susceptibility to αSyn ^A53T^ toxicity or if αSyn ^A53T^ imparts its toxicity by reducing VGLUT levels, we expressed VGLUT RNAi alone or with αSyn ^A53T^. VGLUT knockdown alone had no effect on climbing (Supplementary Figure 4A) or DA neuron loss in females and produced a climbing defect but not DA neuron loss in males (Figure 4A-F). Co-expression of αSyn ^A53T^ with VGLUT RNAi eliminated the sex differences in climbing ability (Supplementary Figure 4A) and DA neuron degeneration (Figure 4A-D) by increasing the severity of αSyn ^A53T^ induced climbing defects and degeneration of PPL1 and PPM1/2 DA neurons in females. Conversely, co-expression of αSyn ^A53T^ with VGLUT RNAi in male DA neurons did not modify the effects of αSyn ^A53T^ on climbing ability (Supplementary Figure 4A) or DA neuron degeneration in PPM1/2 but did increase degeneration in PPL1 (Figure 4A-D). These results suggest that sex differences in climbing and PPM1/2 DA neuron vulnerability to αSyn ^A53T^ are due to differences in VGLUT expression. Interestingly, VGLUT knockdown also sensitized the PPM3 DA neuron cluster to αSyn ^A53T^ toxicity in both males and females leading to degeneration in the previously resistant cluster (Figure 4E-F). These results taken together demonstrate that both sex and region-specific differences in DA neuron vulnerability are mediated by VGLUT expression.

**Figure 4:**
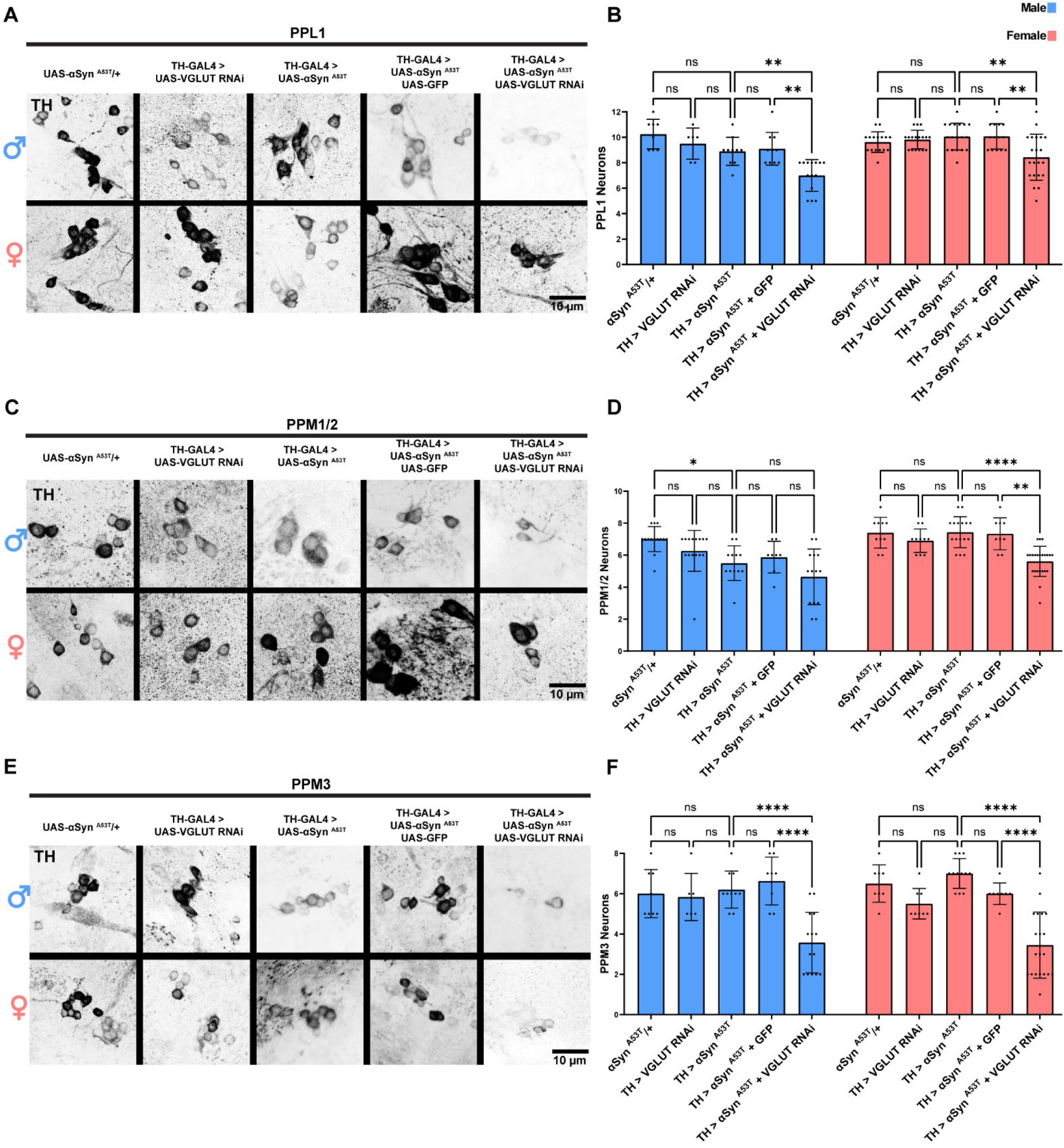
VGLUT knockdown in DA neurons abolishes sex and region-specific differences in vulnerability to *α*Syn ^A53T^. **A** Representative images of male and female PPL1 neurons 35 days post eclosion. **B** Quantification of male and female PPL1 neurons. **C** Representative images of male and female PPM1/2 neurons 35 days post eclosion. **D** Quantification of male and female PPM1/2 neurons. **E** Representative images of male and female PPM3 neurons 35 days post eclosion. **F** Quantification of male and female PPM3 neurons. TH immunofluorescence is black. For all graphs male data is blue and female data is pink. Error bars demonstrate SD. * < 0.05, ** < 0.01, *** < 0.001, **** < 0.0001, N.S = not significant.

### DA neuron VGLUT is required for *α*Syn ^A53T^ induced reduction in mitochondrial number

A multitude of PD linked mutations have been identified in genes encoding proteins that have mitochondrial functions, e.g., PTEN-induced kinase 1 (*PINK1*), Parkin, and DJ-1 ^73–75^. Furthermore, multiple reports have demonstrated that pathogenic αSyn ^A53T^ interacts with mitochondria, impairing function, modifying dynamics/turnover, increasing oxidative stress, and promoting apoptosis via release of cytochrome C ^17–23^. Interestingly, VGLUT has also been linked to mitochondrial function, as knockdown of VGLUT sensitized mitochondria to oxidative stress and altered mitochondrial ATP production ^71^. Based on these reports, we predicted that VGLUT knockdown and αSyn ^A53T^ expression would synergistically affect mitochondrial dynamics. To test this prediction, we used TH-GAL4 to drive MitoTimer (genetically encoded mitochondrial reporter) with UAS-Luciferase RNAi (RNAi against a non-fly mRNA product), UAS-αSyn ^A53T^, UAS-VGLUT RNAi, or UAS-αSyn ^A53T^ with UAS-VGLUT RNAi. Then we assayed both somatic PPL1 cell bodies and synaptic Protocerebral Bridge (PB) mitochondria number, form factor, branch length, and red to green ratio. In PPL1 DA neurons, VGLUT RNAi alone had no effect on mitochondria number in either sex (Figure 5A-B) and reduced both mitochondrial form factor (Figure 5C) and branch length (Supplementary Figure 4B) in males. Conversely, αSyn ^A53T^ expression significantly reduced mitochondria number in both sexes (Figure 5A-B) and reduced both mitochondrial form factor (Figure 5C) and branch length (Supplementary Figure 4B) in females. Co-expression of VGLUT RNAi with αSyn ^A53T^ abolished αSyn ^A53T^ induced reduction of mitochondria number in both sexes (Figure 5A-B) and resulted in a mitochondrial form factor (Figure 5C) and branch length (Supplementary Figure 4B) that was not significantly different from the Luciferase RNAi control for either sex. In the PB, αSyn ^A53T^ reduced mitochondrial number for both sexes (Figure 5D-E) and reduced both mitochondrial form factor (Figure 5F) and branch length (Supplementary Figure 4C) in males. In alignment with our results for PPL1, co-expression of VGLUT RNAi with αSyn ^A53T^ abolished αSyn ^A53T^ induced reduction of mitochondria number in males (Figure 5D-E) and resulted in a mitochondrial form factor (Figure 5F) and branch length (Supplementary Figure 4C) that was not significantly different from the Luciferase RNAi control for either sex. These results demonstrate that VGLUT is required in DA neurons for αSyn ^A53T^ induced changes in mitochondrial number and morphology.

**Figure 5:**
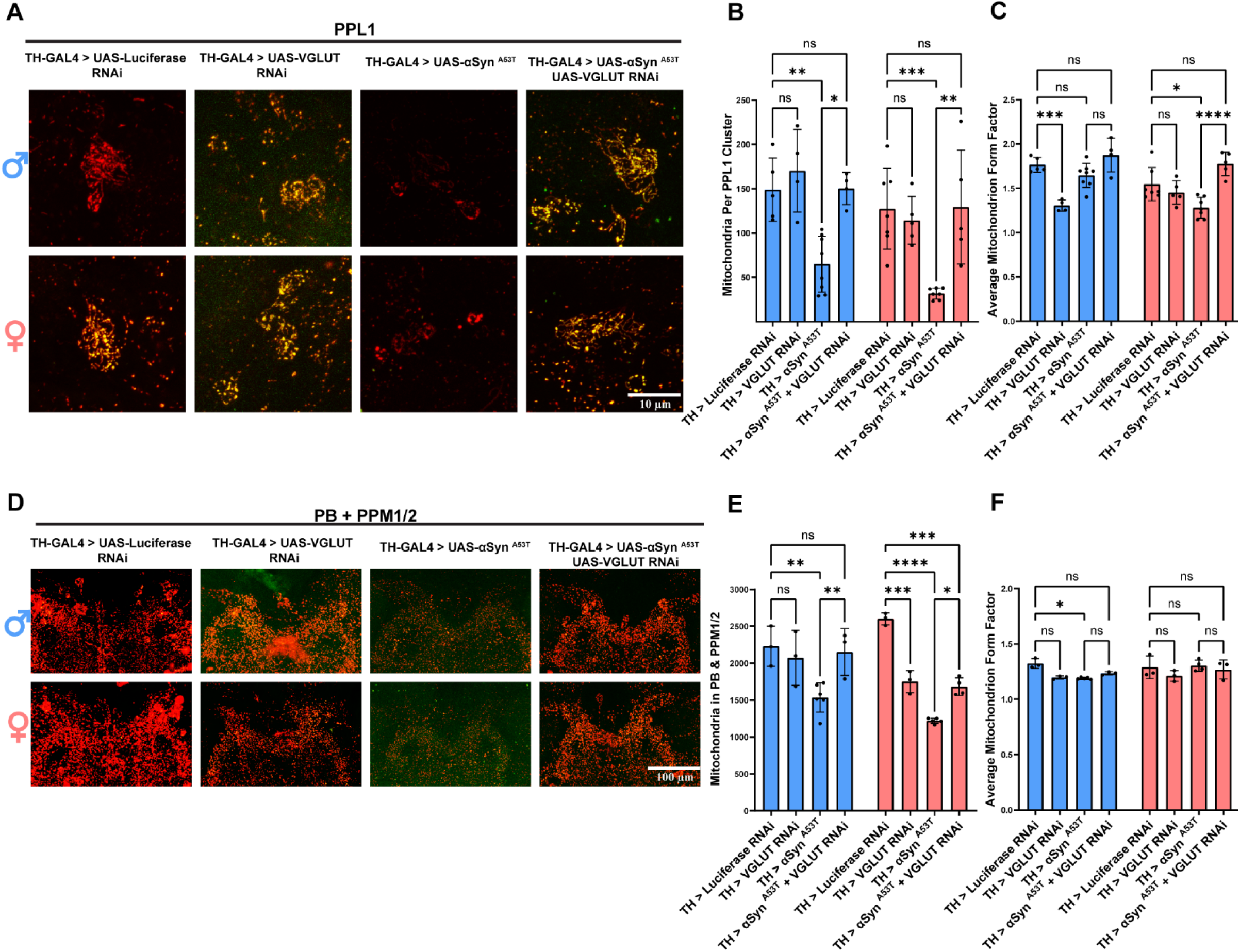
DA neuron VGLUT is required for *α*Syn ^A53T^ induced changes to mitochondrial dynamics. **A** Representative images of PPL1 mitochondria 15 days post eclosion. **B** Quantification of the average number of mitochondria in the PPL1 cluster. **C** Quantification of the average form factor for mitochondria in the PPL1 cluster. **D** Representative images of PB and PPM1/2 mitochondria. **E** Quantification of the average number of mitochondria in the PB and PPM1/2. **F** Quantification of the average form factor for mitochondria in the PB and PPM1/2. For all conditions TH-GAL4 is driving UAS-MitoTimer plus the indicated genotype. For all graphs male data is blue and female data is pink. Error bars demonstrate SD. * < 0.05, ** < 0.01, *** < 0.001, **** < 0.0001, N.S = not significant.

In PPL1, expression of αSyn ^A53T^, VGLUT RNAi, and αSyn ^A53T^ with VGLUT RNAi all reduced the ratio of red to green fluorescence relative to the Luciferase RNAi control (Supplementary Figure 4D). In the PB, αSyn ^A53T^ expression also reduced the ratio of red to green fluorescence relative to the Luciferase RNAi control (Supplementary Figure 4E). This decrease in the ratio of red to green fluorescence was abolished by co-expression VGLUT RNAi (Supplementary Figure 4E). Together with the analysis of mitochondrial number, these results suggest that DA neurons increase mitochondrial turnover in response to αSyn ^A53T^ in a VGLUT dependent manner.

### Increasing total or cytosolic DA phenocopies VGLUT knockdown

VGLUT increases the loading of DA into synaptic vesicles, reducing cytosolic DA levels ^52,53^. This has been proposed, but not yet tested, as the mechanism by which VGLUT protects DA neurons in PD ^9,47,51,53,54^, as cytosolic dopamine is highly reactive and toxic ^24–29^. Our MitoTimer results demonstrate that reducing DA neuron VGLUT levels alters mitochondrial morphology and blocks αSyn ^A53T^ induced mitochondrial turnover. These results are consistent with VGLUT knockdown increasing cytosolic DA, as excess DA can reduce Parkin levels, potentially inhibiting mitochondrial turnover in response to damage ^80^, as well as alter localization of mitochondrial fission and fusion proteins ^80,81^. To test if VGLUT knockdown sensitizes DA neurons to αSyn ^A53T^ pathology by increasing cytosolic DA, we pharmacologically increased total DA levels and genetically increased cytosolic DA levels to determine if these manipulations phenocopied VGLUT knockdown. To increase total DA levels, we aged flies on food containing 10mM L-DOPA. L-DOPA treatment sensitized both male and female DA neurons to αSyn ^A53T^ pathology, increasing αSyn ^A53T^ induced degeneration of PPL1 neurons in males (Figure 6A-B) and PPM1/2 neurons in females (Figure 6C-D). To increase cytosolic DA, we used VMAT RNAi to knockdown VMAT alone or with expression αSyn ^A53T^. Similar to VGLUT knockdown, VMAT knockdown alone had no effect on climbing (Supplementary Figure 5A) or DA neuron loss (Figure 6A-F) in females and produced a climbing defect (Supplementary Figure 5A) but not DA neuron loss (Figure 6A-F) in males. Co-expression of αSyn ^A53T^ with VMAT RNAi decreased female climbing ability (Supplementary Figure 5A) and caused DA neuron degeneration in PPL1, PPM1/2, and PPM3 DA neurons in females (Figure 6A-F). Conversely, co-expression of αSyn ^A53T^ with VMAT RNAi in male DA neurons did not modify the effects of αSyn ^A53T^ on climbing ability (Supplementary Figure 5A) or DA neuron degeneration in PPM1/2 (Figure 6C-D) but did increase degeneration in PPL1(Figure 6A-B) and PPM3 (Figure 6E-F), phenocopying VGLUT knockdown.

**Figure 6:**
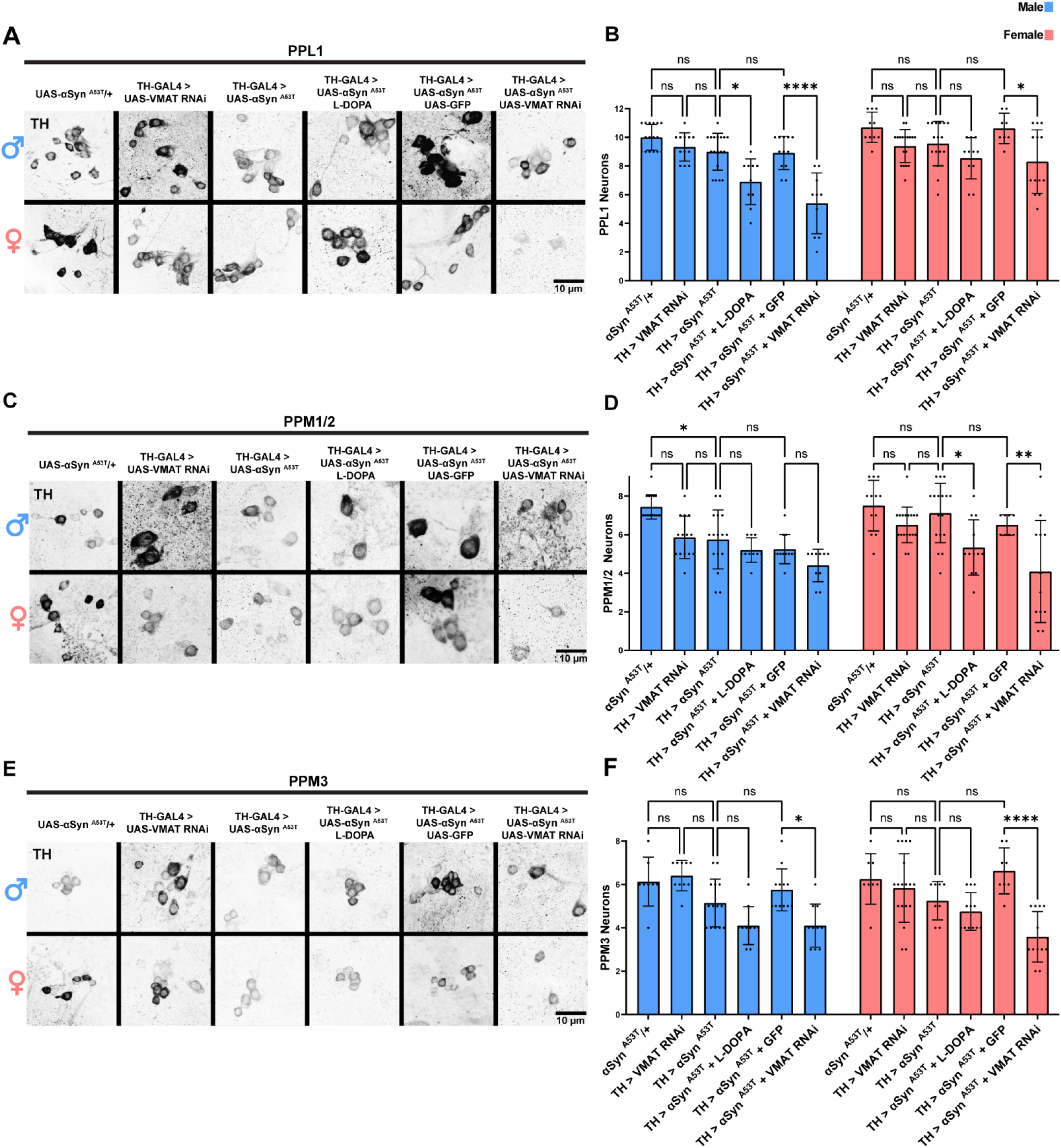
Increasing total or cytosolic DA exacerbates *α*Syn ^A53T^ toxicity. **A** Representative images of male and female PPL1 neurons 35 days post eclosion. **B** Quantification of male and female PPL1 neurons. **C** Representative images of male and female PPM1/2 neurons 35 days post eclosion. **D** Quantification of male and female PPM1/2 neurons. **E** Representative images of male and female PPM3 neurons 35 days post eclosion. **F** Quantification of male and female PPM3 neurons. TH immunofluorescence is black. For all graphs male data is blue and female data is pink. Error bars demonstrate SD. * < 0.05, ** < 0.01, *** < 0.001, **** < 0.0001, N.S = not significant.

To determine if increasing cytosolic DA phenocopies the effects of VGLUT knockdown on DA neuron mitochondria, we used TH-GAL4 to drive MitoTimer with UAS-Luciferase RNAi, UAS-αSyn ^A53T^, UAS-VMAT RNAi, or UAS-αSyn ^A53T^ with UAS-VMAT RNAi. Then we assayed both PPL1 and PB mitochondria number, form factor, branch length, and red to green ratio. In PPL1 DA neurons, VMAT RNAi alone had no effect on mitochondria number (Figure 7A-B), form factor (Figure 7C), or branch length (Supplementary Figure 5B) in either sex. αSyn ^A53T^ expression significantly reduced mitochondria number (Figure 7A-B) and branch length (Supplementary Figure 5B) in both sexes and reduced form factor in females (Figure 7C). Co-expression of VMAT RNAi with αSyn ^A53T^ abolished αSyn ^A53T^’s ability to reduce mitochondria number in both sexes (Figure 7 A-B) and resulted in a mitochondrial form factor in females (Figure 7C) and branch length in males (Supplementary Figure 5B) that was not significantly different from the Luciferase RNAi control. In the PB, αSyn ^A53T^ reduced mitochondrial number for both sexes (Figure 7 D-E) and reduced both mitochondrial form factor (Figure 7F) and branch length (Supplementary Figure 5C) in males. In alignment with our results for PPL1, co-expression of VMAT RNAi with αSyn ^A53T^ abolished αSyn ^A53T^ mediated reduction of mitochondria number in both sexes (Figure 7D-E) and resulted in a mitochondrial form factor (Figure 7F) and branch length (Supplementary Figure 5C) that was not significantly different from the Luciferase RNAi control for either sex. Mirroring our VMAT knockdown results, L-DOPA treatment impaired αSyn ^A53T^ mediated reduction in mitochondrial number for both PPL1 (Supplementary Figure 6A-B) and the PB (Supplementary Figure 6C-D). Like our previous experiment with VGLUT knockdown, in PPL1 expression of αSyn ^A53T^, VMAT RNAi, and αSyn ^A53T^ with VMAT RNAi all reduced the ratio of red to green fluorescence relative to the Luciferase RNAi control (Supplementary Figure 7A). Conversely, in the PB, only αSyn ^A53T^ expression reduced the ratio of red to green fluorescence relative to the Luciferase RNAi control and this decrease was abolished by co-expression of VMAT RNAi ( Supplementary Figure 7B), phenocopying the co-expression of αSyn ^A53T^ with VGLUT RNAi.Together these results demonstrate that increasing total or cytosolic DA levels phenocopy the effects of VGLUT knockdown on locomotor ability, neurodegeneration, and mitochondrial dynamics.

**Figure 7:**
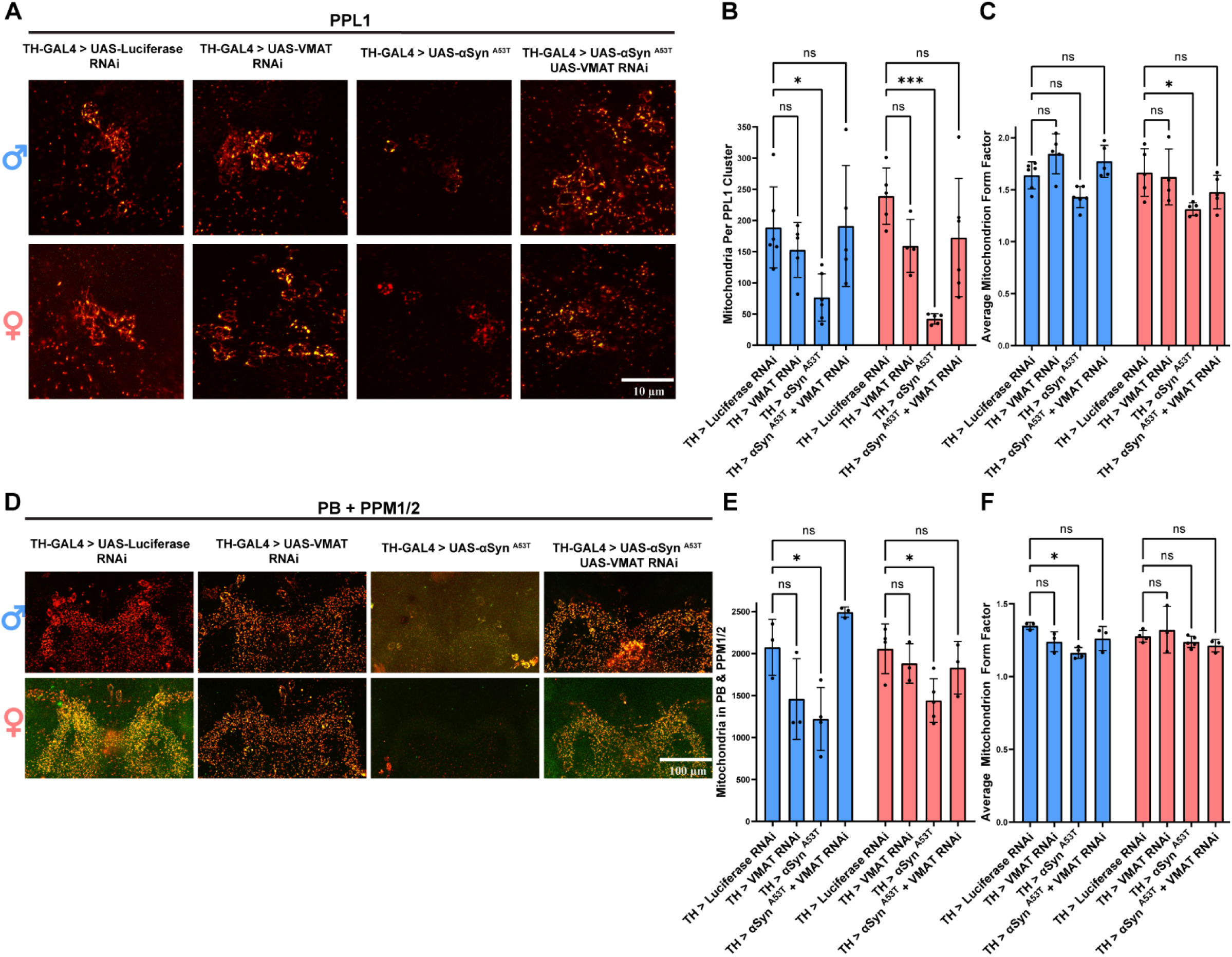
Increasing cytosolic DA phenocopies the effect of VGLUT knockdown on mitochondrial dynamics. **A** Representative images of PPL1 mitochondria 15 days post eclosion. **B** Quantification of the average number of mitochondria in the PPL1 cluster. **C** Quantification of the average form factor for mitochondria in the PPL1 cluster. **D** Representative images of PB and PPM1/2 mitochondria 15 days post eclosion. **E** Quantification of the average number of mitochondria in the PB and PPM1/2. **F** Quantification of the average form factor for mitochondria in the PB and PPM1/2. For all conditions TH-GAL4 is driving UAS-MitoTimer plus the indicated genotype. For all graphs male data is blue and female data is pink. Error bars demonstrate SD. * < 0.05, ** < 0.01, *** < 0.001, **** < 0.0001, N.S = not significant.

### Reducing DA levels partially protects against VGLUT knockdown

Increasing DA phenocopied the effects of VGLUT knockdown, consistent with a model where VGLUT knockdown sensitizes DA neurons to αSyn ^A53T^ pathology by increasing DA levels. However, the presence of these similar phenotypes does not exclude the possibility that an unknown DA independent factor is the cause of the phenotypes produced by VGLUT knockdown. To ascertain whether VGLUT knockdown sensitizes DA neurons by increasing DA levels, we partially inhibited DA synthesis via treatment with the TH inhibitor AMPT and then assessed whether treatment could rescue αSyn ^A53T^ vulnerability induced by VGLUT knockdown. Using TH-GAL4, we co-expressed αSyn ^A53T^ with VGLUT RNAi and aged flies for 35 days in the presence or absence of 15 µM AMPT. Reducing DA levels via AMPT treatment partially rescued αSyn ^A53T^ induced degeneration of male PPM1/2 and female PPM3 DA neurons (Figure 8A–B). These results demonstrate that the increased vulnerability caused by VGLUT knockdown is, at least in part, ameliorated by reducing DA levels.

**Figure 8:**
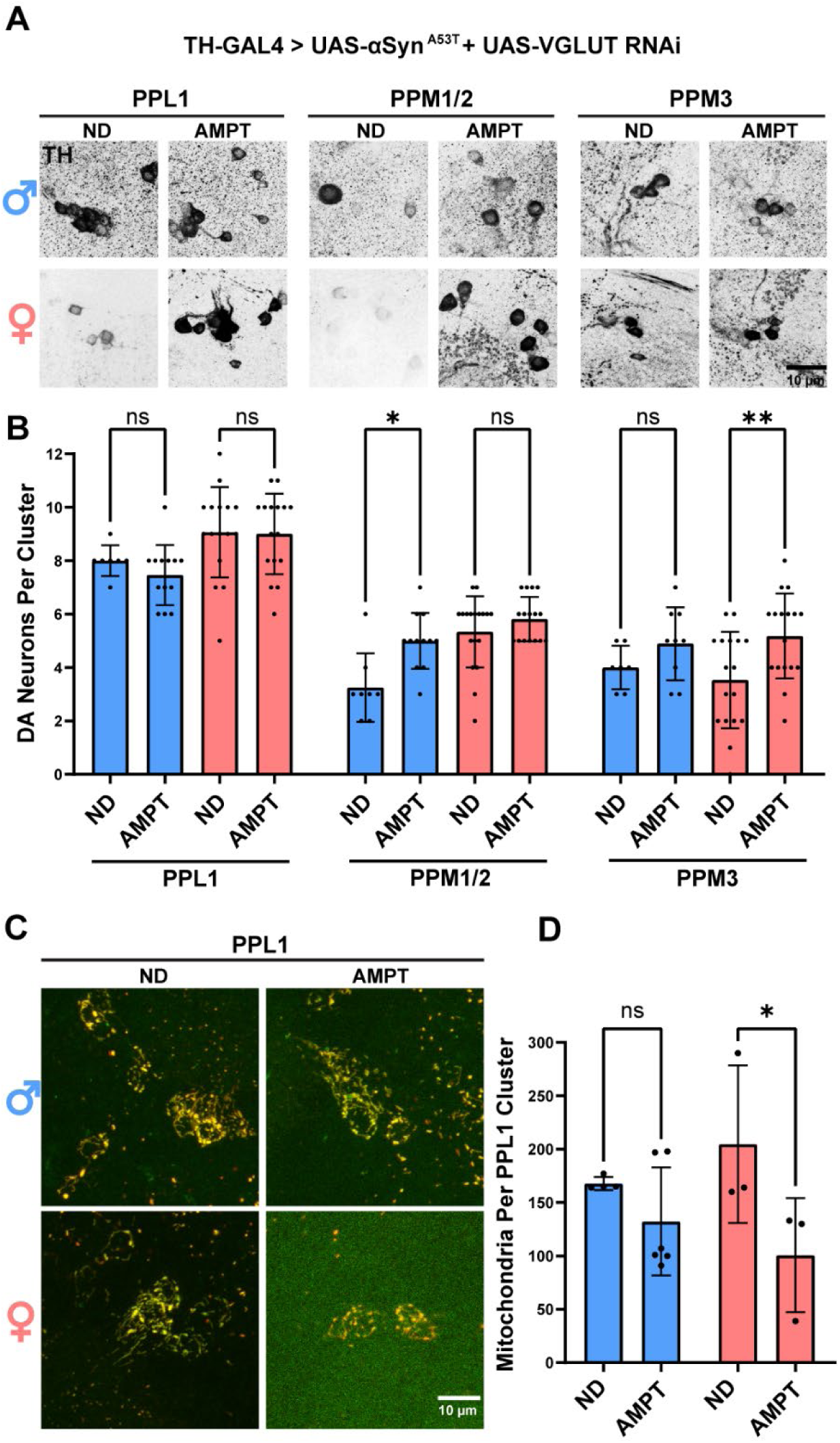
Reducing DA levels partially protects against VGLUT knockdown. **A** Representative images of male and female PPL1, PPM1/2, and PPM3 DA neurons for flies raised in the presence or absence of AMPT. TH immunofluorescence is black. **B** Quantification of male and female DA neurons. **C** Representative images of PPL1 mitochondria 15 days post eclosion for flies raised in the presence or absence of AMPT. **D** Quantification of the average number of mitochondria in the PPL1 cluster. For all graphs male data is blue and female data is pink. ND = No drug. Error bars demonstrate SD. * < 0.05, ** < 0.01, *** < 0.001, **** < 0.0001, N.S = not significant.

To determine if the effects of VGLUT knockdown paired with αSyn ^A53T^ expression on PPL1 and PB DA neuron mitochondria are due to increased DA, we used TH-GAL4 to drive MitoTimer with UAS-αSyn ^A53T^ and UAS-VGLUT RNAi and then aged flies for 15 days in the presence or absence of 15 µM AMPT. AMPT treatment reduced PPL1, but not PB, mitochondria number and form factor in females (Figure 8C-D & Supplementary Figure 8A-F), demonstrating that lowering DA partially blocks the effects of VGLUT knockdown on mitochondrial response to αSyn ^A53T^. These results taken together suggest VGLUT knockdown results in higher levels of cytosolic DA, which leads to DA mediated mitochondrial dysfunction and increased susceptibility to αSyn ^A53T^ toxicity.

## Discussion

Previous studies have demonstrated that VGLUT expressing DA neurons are resistant to degeneration in both postmortem PD human brains and in PD models ^47,54^. However, the mechanism of this protection was unclear. Here, we demonstrate that VGLUT’s protective effect against αSyn ^A53T^ pathology is in part mediated by its ability to reduce cytosolic DA, which allows DA neurons to adjust their mitochondrial dynamics in response to αSyn ^A53T^ expression. Somewhat paradoxically, these adjustments entail decreasing mitochondria number, which likely reduces energy production, and increasing fragmentation of the mitochondrial network, which is typically associated with disease and apoptosis ^82^. In line with our findings, previous studies have demonstrated αSyn ^A53T^ decreases mitochondria number by increasing mitophagy, but it was unclear whether this increase in mitophagy is a protective change or pathological ^19,35,83,84^. Our data suggests that these changes to the mitochondrial network are compensatory, likely clearing damaged mitochondria to prevent cytochrome c leakage and ROS production. In support of this, increasing expression of the mitophagy proteins PINK1 and Parkin protect against αSyn pathology; furthermore, knockout of PINK1or Parkin increases vulnerability to αSyn pathology, likely by interfering with mitophagy ^85–88^.

Our data demonstrates that VGLUT in DA neurons is required for mitochondrial network adaptation to αSyn ^A53T^ because it limits cytosolic DA levels. In vitro reports have demonstrated that DA can affect mitochondrial fission-fusion dynamics by altering localization of Dynamin-Related Protein 1 (DRP1) and levels of Optic Atrophy Type 1 (OPA1) ^80,81^. Tight regulation of mitochondrial fission-fusion dynamics is paramount for the efficient clearance of damaged mitochondria, as fragmented mitochondria are more efficiently cleared via mitophagy ^89–92^. A previous study in *Drosophila* demonstrated that mis-localization of DRP1 can exacerbate αSyn ^A53T^ induced climbing defects and neurodegeneration ^17^. Furthermore, the same study showed that increasing mitochondrial fission via DRP1 overexpression rescued both αSyn ^A53T^ induced climbing defects and neurodegeneration ^17^. Interestingly, co-expression of αSyn ^A53T^ and DRP1 decreased the ratio of MitoTimer red to green fluorescence, consistent with DRP1 protecting by increasing mitochondrial turnover ^17^. DA can also reduce levels of Parkin, which, in addition to its direct role in mitophagy, can indirectly affect clearance by tagging Mitofusins for degradation, thereby promoting mitochondrial fragmentation and facilitating mitochondrial turnover ^80,89^.

Furthermore, excess DA can also cause lysosomal dysfunction, which may further inhibit turnover of damaged mitochondria as well as degradation of αSyn ^26^. This may explain why some previous studies have reported reduced TH immunoreactivity following αSyn ^A53T^ expression, as this reduction may represent a compensatory mechanism to lower DA levels in response to αSyn ^A53T^ pathology ^93,94^.

In our experiments, AMPT treatment only partially rescued VGLUT knockdown. This could be because a higher concentration is needed for a complete rescue or because VGLUT protects against αSyn ^A53T^ pathology by an additional mechanism other than just increasing DA loading. A previous report demonstrated that VGLUT2 knockout reduces brain derived neurotrophic factor (BDNF) and its receptor tropomyosin receptor kinase B (TrkB) expression in DA neurons ^95^. BDNF is neuroprotective and has been tested as a potential therapeutic agent for PD; thus, VGLUT/VGLUT2 may also protect DA neurons by regulating BDNF and TrkB levels ^96^. Another potential mechanism might be that VGLUT expression during development promotes expression of glutamatergic neurotransmission machinery, resulting in increased intracellular glutamate availability. Glutamate has been demonstrated to protect against DA auto-oxidation, resulting in decreased levels of reactive DA species and ROS ^97^. Furthermore, glutamate can be converted to the antioxidant glutathione, which is upregulated in response to increased DA levels and can protect against DA induced apoptosis ^47,98,99^.

Our results demonstrated that there are sex differences in *Drosophila* DA neuron vulnerability to αSyn ^A53T^ pathology and, by selectivity masculinizing female DA neurons, showed that these differences are cell autonomous. Females have higher VGLUT in DA neurons and VGLUT knockdown in DA neurons abolishes these sex differences, demonstrating that the cause of the sex difference is differential VGLUT expression. Two recent papers demonstrated that pan neuronal expression of αSyn ^A53T^ decreases median lifespan in male flies more than in female flies, suggesting that other non-dopaminergic factors may also protect females against αSyn ^A53T^ pathology ^64,65^. Furthermore, a recent report using mice expressing αSyn ^A53T^ and inoculated with recombinant human αSyn preformed fibrils reported more aggressive neurodegeneration in males than in females, affecting both dopaminergic and non-dopaminergic regions ^100^. Therefore, more work needs to be done to fully elucidate all the factors that protect females against αSyn ^A53T^ pathology.

In this report and previous reports, PPM3 neurons have been shown to be resistant to αSyn induced degeneration ^93,101,102^. These neurons have also been shown to resist degeneration and mitochondrial dysfunction in other PD models, suggesting they are generally resistant to DA neuron stressors ^76,103–105^. Our data suggest that this increased resilience is in part due to VGLUT expression, as VGLUT knockdown makes them extremely sensitive to αSyn ^A53T^ pathology. Moderately increasing VGLUT expression in other DA neuron clusters might theoretically increase their resilience as well; however, VGLUT overexpression is toxic and can cause severe neurodegeneration via excitotoxicity^106^. Interestingly, this toxicity also extends to the VGLUT expressing neuron itself, suggesting an additional cell-autonomous mechanism of VGLUT mediated cell death ^107^. The mechanism of this cell-autonomous cell death is unknown but may be due to VGLUT’s role as a phosphate transporter, as VGLUT can increase intracellular levels of phosphate when overexpressed or during times of high activity ^108–111^. High intracellular phosphate levels can be toxic, causing apoptotic cell death ^112,113^.

## Acknowledgments

The authors thank Dr. Hermann Aberle for donating the VGLUT antibody and Dr. Gerald Rubin for providing the UAS-GFP stock used in this study. We also thank Tyler Marquardt for computer access, which was used for figure generation and data analysis, as well as Stefan Choy, Dominick Costanzo, and Joshua November for feedback on the manuscript. This study was supported by a grant from the NIH (R03NS144936) to DTB.

---

**Supplementary Figure 1:**
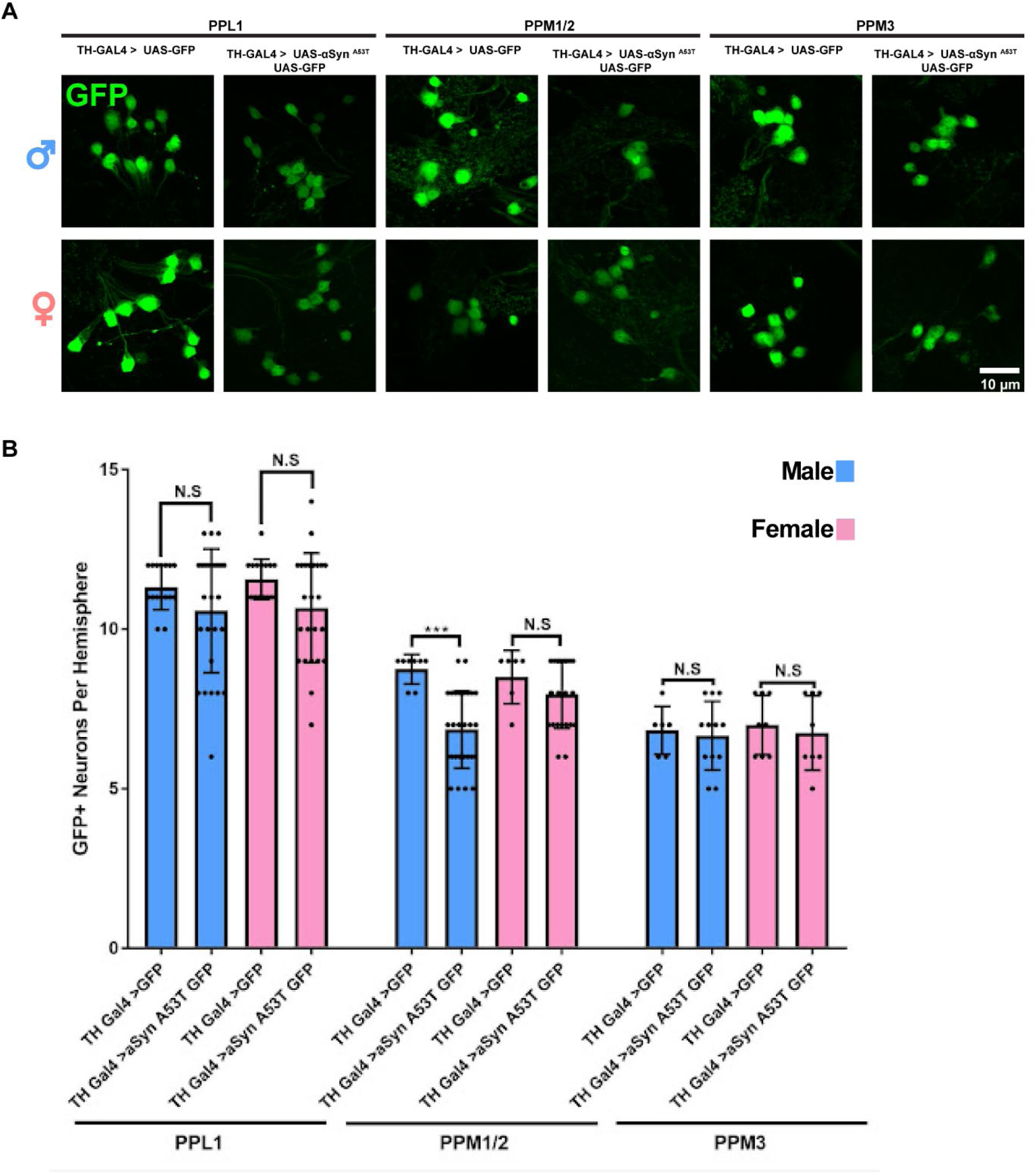
Male flies exhibit selective DA neuron degeneration in response to *α*Syn ^A53T^ expression. **A** Representative images of male and female PPL1, PPM1/2, and PPM3 DA neurons. GFP immunofluorescence is green. **B** Quantification of male and female DA neurons 35 days post eclosion. Error bars demonstrate SD. * < 0.05, ** < 0.01, *** < 0.001, **** < 0.0001, N.S = not significant.

**Supplementary Figure 2:**
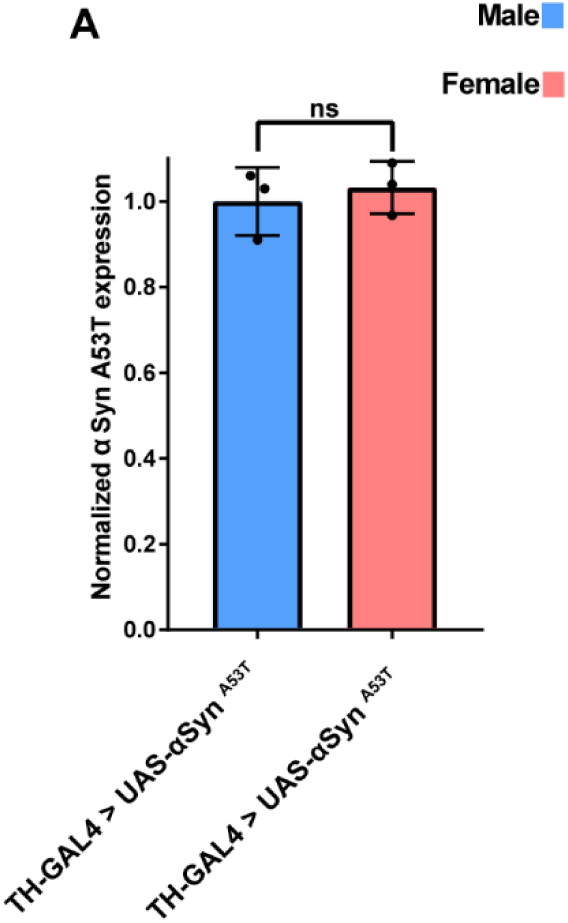
Sex differences are not due to differential transgene expression. A Quantification of male and female *α*Syn ^A53T^ expression. Male data is blue and female data is pink. Error bars demonstrate SD. * < 0.05, ** < 0.01, *** < 0.001, **** < 0.0001, N.S = not significant.

**Supplementary Figure 3:**
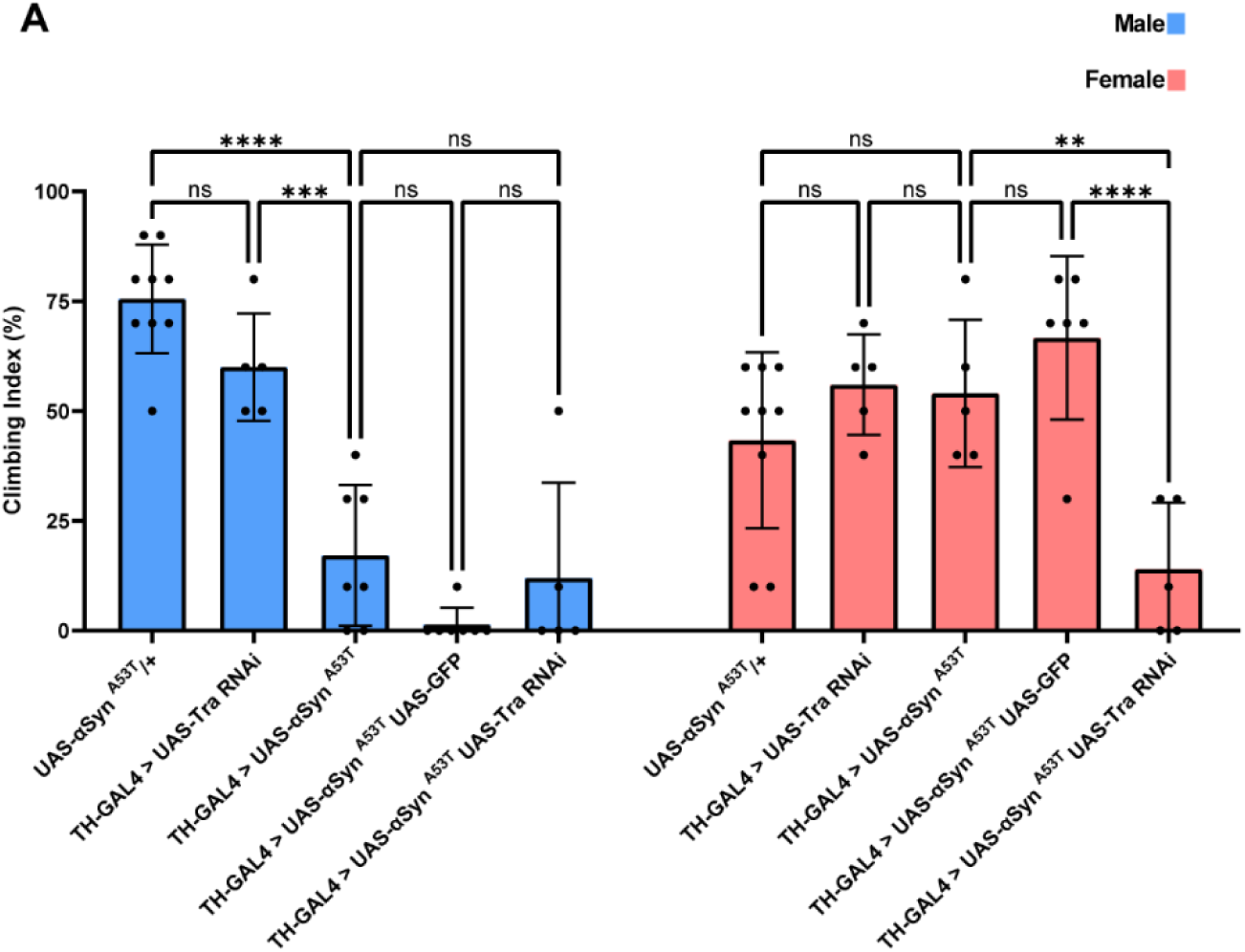
Sex differences in *α*Syn ^A53T^ induced climbing defects are cell autonomous. **A** Climbing assay data for male and female flies 35 days post eclosion. Each point on the graph represents a vial of 10 flies. Male data is blue and female data is pink. Error bars demonstrate SD. * < 0.05, ** < 0.01, *** < 0.001, **** < 0.0001, N.S = not significant.

**Supplementary Figure 4:**
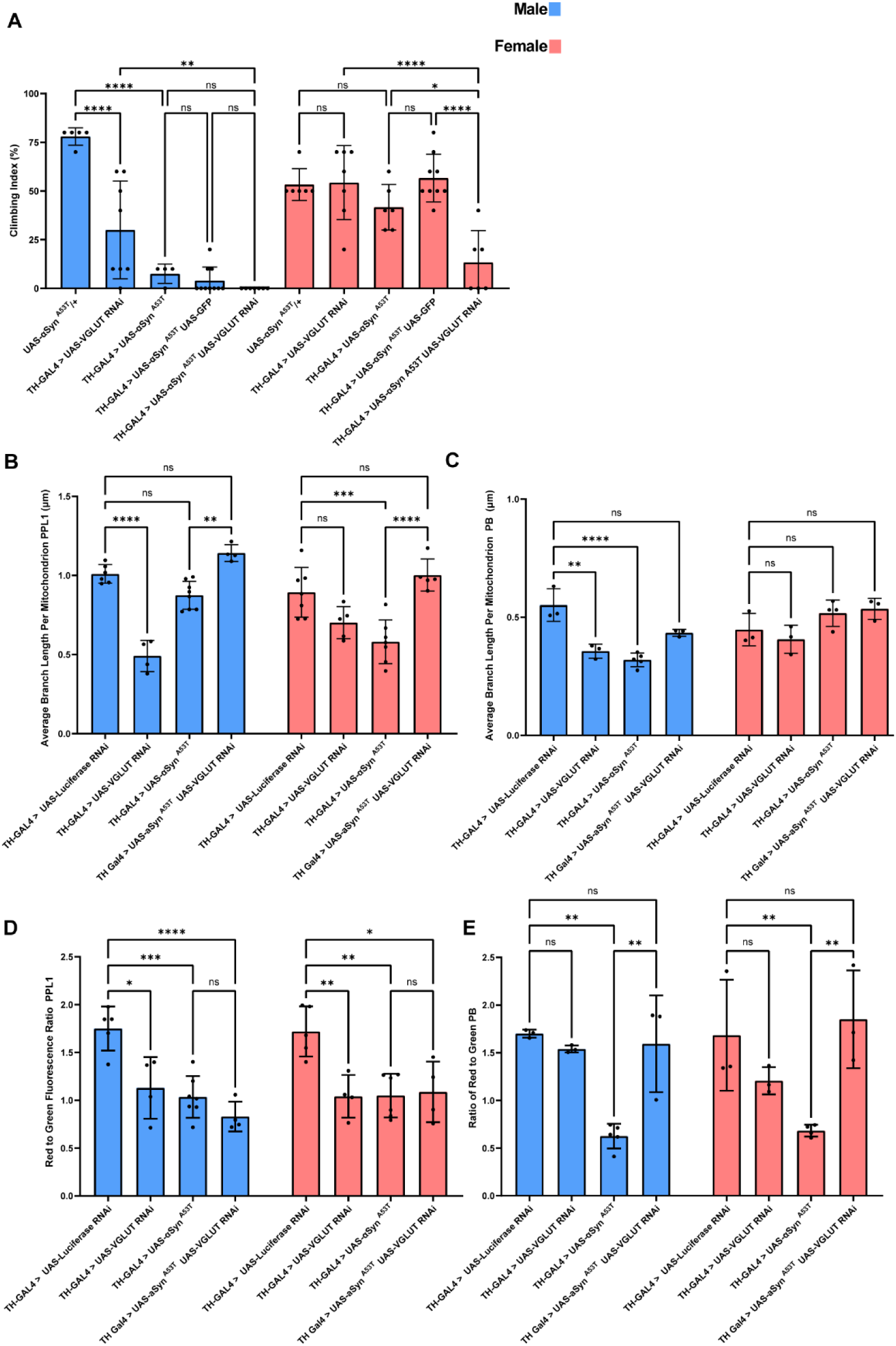
DA neuron knockdown of VGLUT abolishes sex differences in *α*Syn ^A53T^ induced climbing defects and mitochondrial dynamics. **A** Climbing assay data for male and female flies 35 days post eclosion. Each point on the graph represents a vial of 10 flies. **B** Quantification of the average branch length for mitochondria in the PPL1 cluster. **C** Quantification of the average branch length for mitochondria in the PB and PPM1/2 cluster. **D** Quantification of the ratio of red to green fluorescence for mitochondria in PPL1. **E** Quantification of the ratio of red to green fluorescence for mitochondria in the PB and PPM1/2. For all conditions in B-E TH-GAL4 is driving UAS-MitoTimer plus the indicated genotype. Male data is blue and female data is pink. Error bars demonstrate SD. * < 0.05, ** < 0.01, *** < 0.001, **** < 0.0001, N.S = not significant.

**Supplementary Figure 5:**
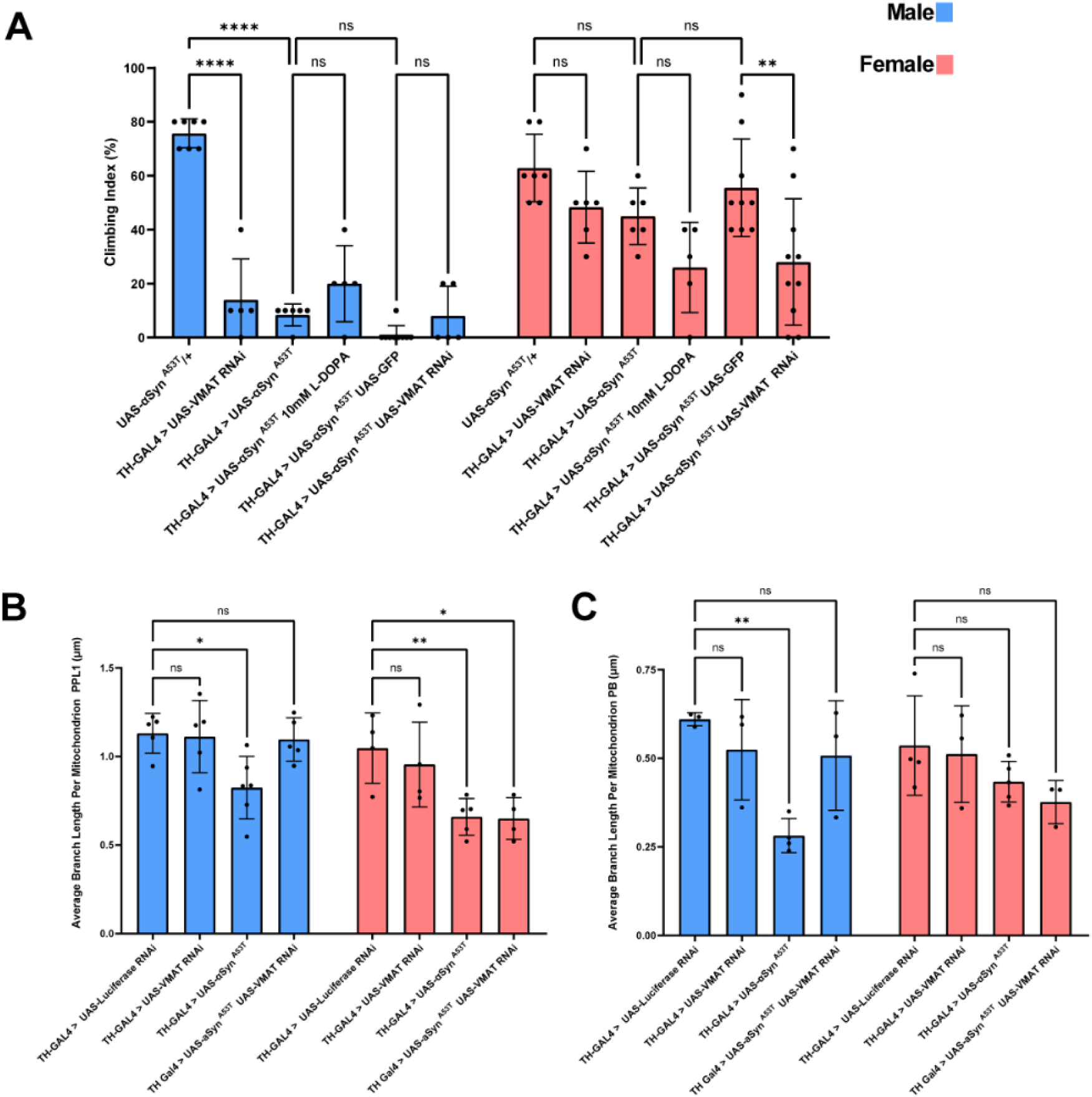
VMAT knockdown in DA neurons mimics the effects of VGLUT knockdown on *α*Syn ^A53T^ induced climbing defects and mitochondrial dynamics. **A** Climbing assay data for male and female flies 35 days post eclosion. Each point on the graph represents a vial of 10 flies. **B** Quantification of the average branch length for mitochondria in the PPL1 cluster. **C** Quantification of the average branch length for mitochondria in the PB and PPM1/2 cluster. For all conditions in B-C TH-GAL4 is driving UAS-MitoTimer plus the indicated genotype. Male data is blue and female data is pink. Error bars demonstrate SD. * < 0.05, ** < 0.01, *** < 0.001, **** < 0.0001, N.S = not significant.

**Supplementary Figure 6:**
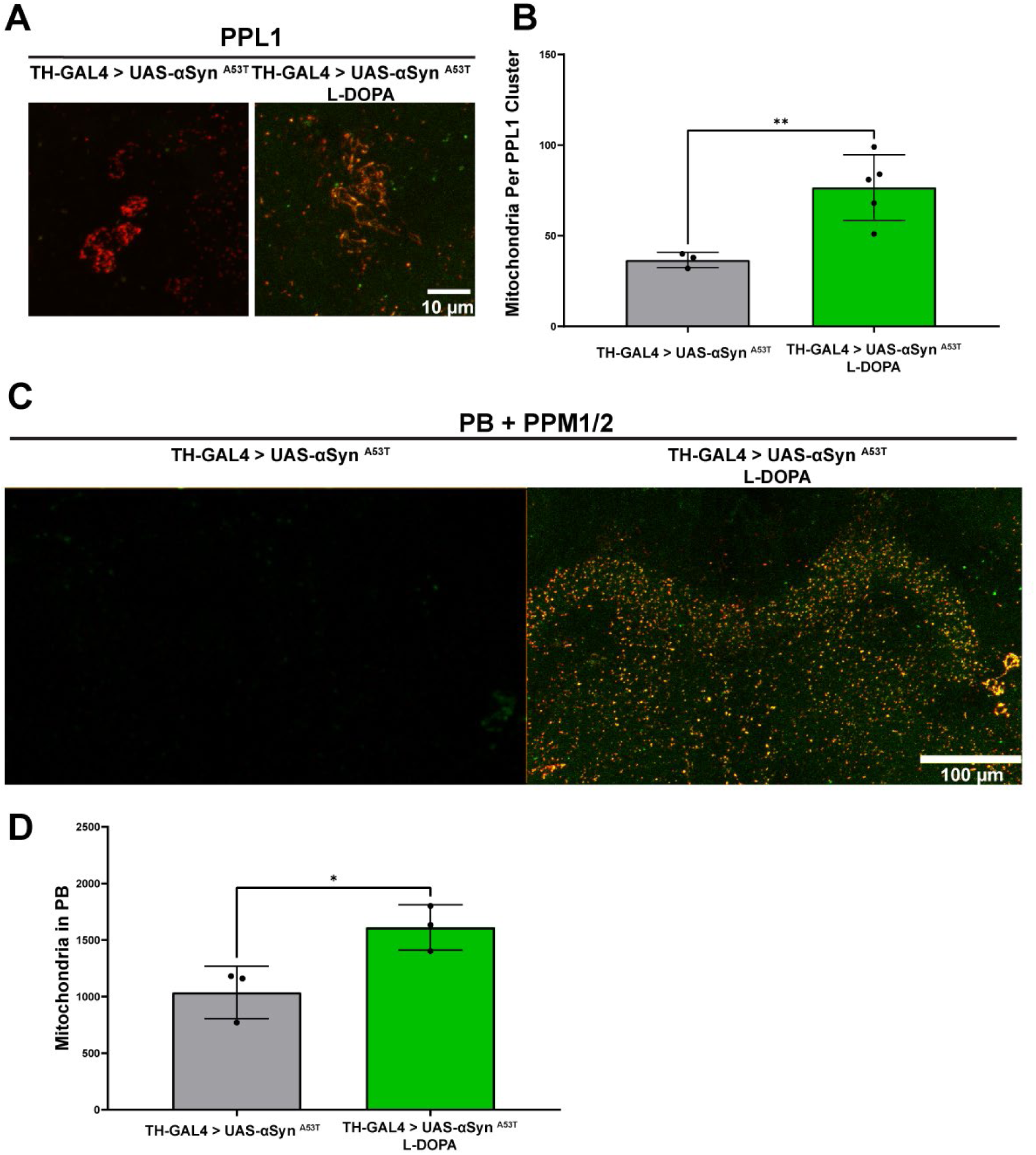
Increasing total DA phenocopies the effect of VGLUT knockdown on mitochondrial number. **A** Representative images of PPL1 mitochondria 15 days post eclosion. **B** Quantification of the average number of mitochondria in the PPL1 cluster. **C** Representative images of PB and PPM1/2 mitochondria 15 days post eclosion. **D** Quantification of the average number of mitochondria in the PB and PPM1/2. For all conditions TH-GAL4 is driving UAS-MitoTimer plus the indicated genotype. Error bars demonstrate SD. * < 0.05, ** < 0.01, *** < 0.001, **** < 0.0001, N.S = not significant.

**Supplementary Figure 7:**
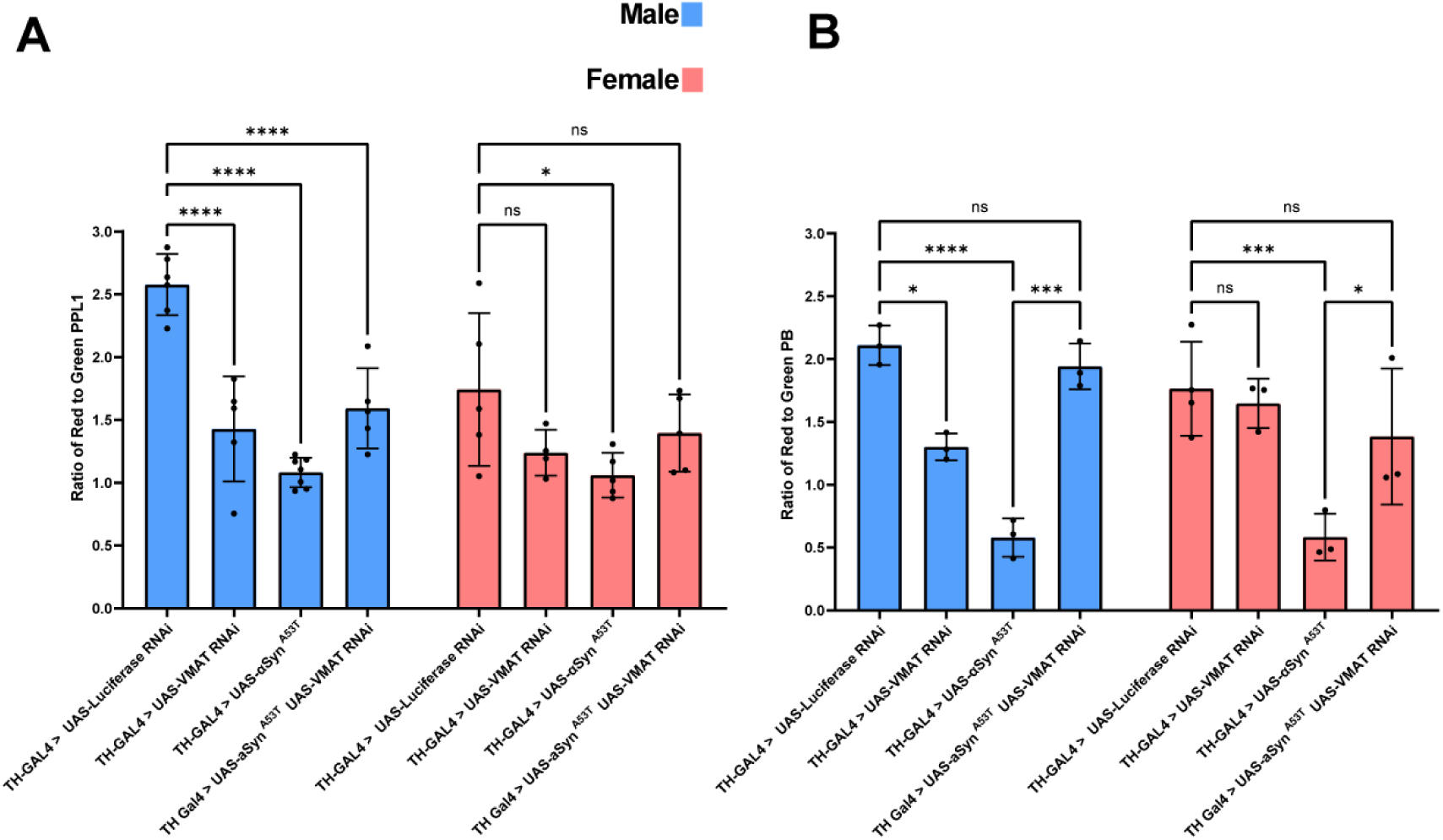
VMAT knockdown in DA neurons mimics the effects of VGLUT knockdown on *α*Syn ^A53T^ induced mitochondrial turnover. **A** Quantification of the ratio of red to green fluorescence for mitochondria in PPL1. **B** Quantification of the ratio of red to green fluorescence for mitochondria in the PB and PPM1/2. For all conditions TH-GAL4 is driving UAS-MitoTimer plus the indicated genotype. Male data is blue and female data is pink. Error bars demonstrate SD. * < 0.05, ** < 0.01, *** < 0.001, **** < 0.0001, N.S = not significant.

**Supplementary Figure 8:**
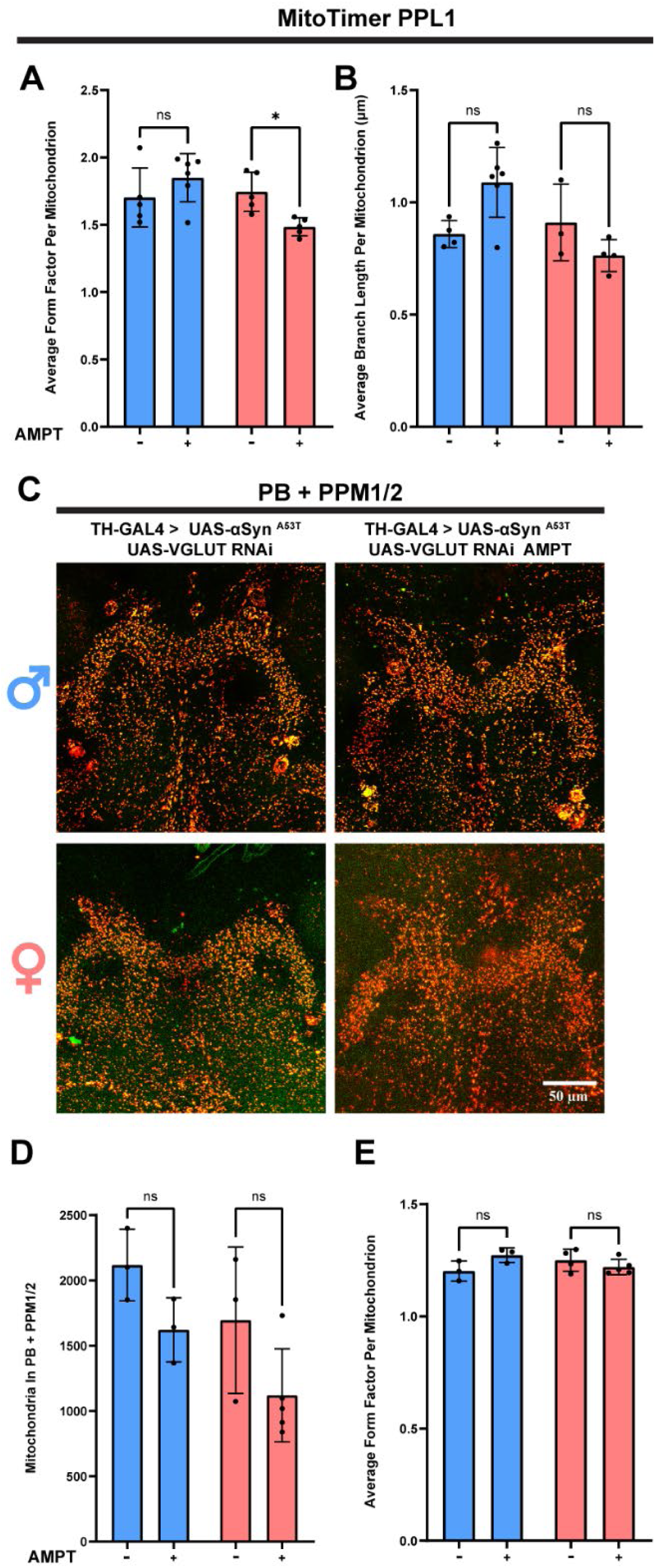
Reducing DA levels partially protects against the effects of VGLUT knockdown on mitochondria. **A** Quantification of the average form factor for mitochondria in the PPL1 cluster. **B** Quantification of the average branch length for mitochondria in the PPL1 cluster. **C** Representative images of PB and PPM1/2 mitochondria. **D** Quantification of the average number of mitochondria in the PB and PPM1/2. **E** Quantification of the average form factor for mitochondria in the PB and PPM1/2. **+** = AMPT treatment. For all conditions TH-GAL4 is driving UAS-MitoTimer plus the indicated genotype. For all graphs male data is blue and female data is pink. Error bars demonstrate SD. * < 0.05, ** < 0.01, *** < 0.001, **** < 0.0001, N.S = not significant

## Notes

### Competing Interest Statement

The authors have declared no competing interest.

